# Identification of *Klebsiella* capsule synthesis loci from whole genome data

**DOI:** 10.1101/071415

**Authors:** Kelly L. Wyres, Ryan R. Wick, Claire Gorrie, Adam Jenney, Rainer Follador, Nicholas R. Thomson, Kathryn E. Holt

## Abstract

**Background:** *Klebsiella pneumoniae* and close relatives are a growing cause of healthcare-associated infections for which increasing rates of multi-drug resistance are a major concern. The *Klebsiella* polysaccharide capsule is a major virulence determinant and epidemiological marker. However, little is known about capsule epidemiology since serological typing is not widely accessible, and many isolates are serologically non-typeable. Molecular methods for capsular typing are needed, but existing methods lack sensitivity and specificity and fail to take advantage of the information available in whole-genome sequence data, which is increasingly being generated for surveillance and investigation of *Klebsiella*.

**Methods:** We investigated the diversity of capsule synthesis loci (K loci) among a large, diverse collection of 2503 genome sequences of *K. pneumoniae* and closely related species. We incorporated analyses of both full-length K locus DNA sequences and clustered protein coding sequences to identify, annotate and compare K locus structures, and we propose a novel method for identifying K loci based on full locus information extracted from whole genome sequences.

**Results:** A total of 134 distinct K loci were identified, including 31 novel types. Comparative analysis of K locus gene content detected 508 unique protein coding gene clusters that appear to reassort via homologous recombination, generating novel K locus types. Extensive nucleotide diversity was detected among the *wzi* and *wzc* genes, both within and between K loci, indicating that current typing schemes based on these genes are inadequate. As a solution, we introduce *Kaptive*, a novel software tool that automates the process of identifying K loci from large sets of *Klebsiella* genomes based on full locus information.

**Conclusions:** This work highlights the extensive diversity of *Klebsiella* K loci and the proteins that they encode. We propose a standardised K locus nomenclature for *Klebsiella*, present a curated reference database of all known K loci, and introduce a tool for identifying K loci from genome data (https://github.com/katholt/Kaptive). These developments constitute important new resources for the *Klebsiella* community for use in genomic surveillance and epidemiology.

## Background

*Klebsiella pneumoniae, Klebsiella variicola* and *Klebsiella quasipneumoniae* are ubiquitous, encapsulated Gram-negative bacteria. They can be carried asymptomatically in the human gut or nasopharynx [1] but are also opportunistic pathogens, frequently associated with human disease and recognised as a significant threat to global health. Antimicrobial resistance, particularly multi-drug resistance and resistance to the carbapenems, is a major concern. Notably, there are a number of multi-drug resistant clones, which are distributed world-wide and are reported to cause outbreaks of healthcare-associated infections [2,3]. There are also increasing reports of invasive, community-acquired *K. pneumoniae* disease in many Asian countries [4]. While this phenomenon is not yet well understood, it is associated with ‘hypervirulent’ *K. pneumoniae* strains expressing specific capsular serotypes known as K1, K2 and K5 [5,6].

In order to control the emerging threat of *K. pneumoniae sensu stricto, K. variicola* and *K. quasipneumoniae* (hereafter collectively referred to as *K. pneumoniae* unless otherwise stated), there is an urgent requirement for genome-based surveillance. Recent advances in understanding *K. pneumoniae* population structure [7,8] highlight immense genomic diversity and provide a framework for tracking this pathogen. Useful strategies involve analyses of lineages or multi-locus sequence types in combination with resistance and virulence gene characterisation, e.g. using tools such as SRST2 [9] and BIGSdb [8]. Additionally, there have been successful *K. pneumoniae* outbreak investigations using genomic analysis [3,10]. However, methods for tracking *K. pneumoniae* capsular variation are currently lacking.

The polysaccharide capsule is the outer most layer of the *K. pneumoniae* cell, which protects the bacterium from desiccation, phage and protist predation [11]. The capsule is also a key virulence determinant due to its antiphagocytic properties [1214]. There are 77 immunologically distinct *K. pneumoniae* capsule types (K-types) defined by serology, mostly based on work done in the 1950s-70s [15–17]. However, serological typing requires specialist techniques and reagents not available to most microbiology laboratories, so it is very rarely applied. Furthermore, between 10% and 70% of *K. pneumoniae* isolates are serologically non-typeable, either because they express a novel capsule (most commonly for clinical isolates) or are noncapsulated [6,18,19].

*K. pneumoniae* employ a Wzy-dependent capsule synthesis process [11,20] and the genes required for capsule synthesis and assembly are located at the capsule polysaccharide synthesis locus (K locus). The K locus varies in length from 10 to 30 kbp [21–26] and includes a set of ‘common’ genes in the terminal regions, which encode the core capsule biosynthesis machinery (e.g. *galF, wzi, wza, wzb, wzc, gnd* and *ugd*). The central region of the K locus is highly variable, encoding the capsule-specific sugar synthesis, processing and export proteins plus the core assembly components Wzx (flippase) and Wzy (capsule repeat unit polymerase). The *wzx* and *wzy* genes are more diverse than those of the other core assembly components and do not have a fixed position within the K locus [22].

K locus nucleotide sequences and annotations are now available for a large number of *K. pneumoniae* isolates, including the 77 K-type reference strains [3,21–23,25,27–29]. Serological K-types are generally defined by distinct sets of genes in the variable central region of the K locus. This is usually due to the presence of entirely different sets of protein coding sequences; however two types (K22 and K37) are distinguished by a single point mutation resulting in a premature stop codon that affects acetyltransferase function [22].

A number of molecular K-typing schemes have been developed that take advantage of the conserved K locus structure: restriction fragment length polymorphism Retyping’) [30], *wzi* and *wzc* typing [31,32] and capsule-specific *wzy* PCR-based typing [25,33]. C-typing comprises PCR amplification of a large region of the K locus (from upstream of *wzi* to within *gnd*), followed by *HincII* restriction. In contrast, *wzi* and *wzc* typing each comprise PCR amplification and nucleotide sequencing of regions of a single gene, *wzi* and *wzc* respectively. Within the *wzi* scheme, unique alleles are associated to specific K-types [32]. Within the *wzc* scheme, K-types are assigned based on the level of *wzc* nucleotide similarity to a reference sequence, with a threshold of 94% [31]. These molecular typing methods are less technically challenging than serological techniques and are more discriminatory [30–32]. However, none of the methods have been widely adopted and regardless of the method, a substantial proportion of isolates remain non-typeable. As a consequence, the true extent of *K. pneumoniae* capsule diversity remains unknown.

Here we report the K loci from a collection of 2503 *K. pneumoniae*. We identify 31 novel K loci, and provide evidence that limited additional diversity remains to be discovered in *K. pneumoniae*. We define a standardised nomenclature for *Klebsiella* K loci, provide a curated reference database and introduce *Kaptive*, a tool for rapid identification of reference K loci from genome data, which will greatly facilitate genomic surveillance efforts and evolutionary investigations of this important pathogen.

## Methods

We obtained a total of 2600 *K. pneumoniae* genomes (2021 publicly available genomes and 579 novel genomes from a diverse set of isolates collected in Australia). Sequence reads were generated locally or obtained from the European Nucleotide Archive (accessions listed in **Table S1, Additional File 1**); 916 genomes that were publicly available as assembled contigs only were downloaded from PATRIC [34] and the NCTC3000 project [35]. For isolates sequenced in this study (n = 579) DNA was extracted and libraries prepared using the Nextera^®^ XT 96 barcode DNA kit and 125 bp paired-end sequence reads were generated on the Illumina HiSeq 2500 platform.

All paired-end read sets were filtered to remove reads with mean Phred quality score <30, then assembled *de novo* using SPAdes v3 [36]. Genomes were excluded from the study if they were duplicate samples, if there was evidence of contamination or mixed culture measured by: (i) <50% reads mapped to the NTUH-K2044 reference chromosome (accession: AP006725.1); (ii) the ratio of heterozygous/homozygous SNP calls compared to the reference chromosome exceeded 20% ; (iii) the total assembly length was >6.5Mb, or >6.0Mb with evidence of >1% *non-Klebsiella* read contamination as determined by MetaPhlAn [37]; or (v) the assembly was low quality i.e total length <5Mb.

### Existing high quality K locus reference sequences

K locus nucleotide sequences for each of the 77 K-type references and two serologically non-typeable strains published previously [21–26] were obtained from GenBank or directly from the authors (accessions in **Table S2, Additional File 2**). A total of 12 additional K locus sequences had been published prior to the K-type references [3,27,29,38]; we have compared these loci to those of the 77 K-type references [21–26] and identified seven that were novel. These seven novel loci plus 17 distinct loci described in our recent survey [28] were added to the non redundant list of K locus reference sequences, resulting in a total of 103 loci (see **Table S2, Additional File 2**).

### Identification of novel K loci

In order to identify novel K loci we first classified each genome by similarity to previously known loci. BLASTn [39] was used to search each genome assembly for sequences with similarity to those of annotated K locus coding sequences (CDS) usually located between *galF* and *ugd* inclusive (minimum coverage 80%, minimum identity 50%). Transposase CDS present in the published K locus reference sequences were excluded from this analysis since they are not K locus specific. Up to three missing CDS were tolerated for K locus assignment, to allow for assembly problems and insertion sequence (IS) insertions (see **Figure S1, Additional File 3**). This approach can successfully distinguish the 77 K-type reference loci (with the exception of K22 and K37).

Genomes that could not be assigned a K locus were investigated further: BLASTn was used to identify the *galF* and *ugd* genes within the assembly, and single contig loci were extracted. The assembly graph viewer Bandage [40] was used to identify K loci that did not assemble on a single contig or where *galF* and/or *ugd* where missing. Loci were clustered, with identity and coverage thresholds of 90%, using CD-HIT-EST [41,42]. A representative sequence from each cluster was annotated with Prokka [43], using all proteins in the 77 reference K-type loci as the primary annotation database. Novel K locus sequences were deposited in Genbank (accessions: LT603702 - LT603735; also included in the Kaptive database at https://github.com/katholt/Kaptive/).

Recombination in K loci was investigated by aligning nucleotide sequences for the eight common genes (extracted from the reference annotations) using MUSCLE [44]. This generated a 9944 bp concatenated gene alignment which was used as input for maximum likelihood (ML) phylogenetic inference with RAxML [45] (best scoring tree from five runs each of 1000 bootstrap replicates with gamma model of rate heterogeneity), and recombination analysis using ClonalFrameML [46] (run for 1000 simulations and using the ML phylogeny as the starting tree).

### Amino acid clustering

Predicted amino acid sequences of all annotated K locus coding regions were translated from the DNA sequences using BioPython and clustered with CD-HIT [41,42] (90%, 80%, 70%, 60%, 50%, 40% identity). We explored the co-occurrence of predicted protein clusters present in three or more K loci each (excluding the common proteins and the initiating glycosyltransferases, WbaP and WcaJ, n = 115 clusters for analysis): Pairwise Jaccard similarity scores were calculated as *J (A,B*) = *A* ∩ *B* / *A* U *B* and were used to draw a weighted edge graph with the igraph R package v 1.0.1 [47]. A weight threshold was determined empirically as 0.61 and all edges for which *J <* 0.61 were removed.

### Wzc and wzi nucleotide sequence determination

We characterised *wzi* and *wzc* sequence diversity and explored the association with K loci. We used SRST2 [9] to determine *wzi* alleles defined in the *K. pneumoniae* BIGSdb [48]. BLASTn was used to determine alleles in genomes available only as assemblies. Novel alleles were submitted to the *K. pneumoniae* BIGSdb for official designation. *Wzc* sequences were extracted from genome assemblies by BLASTn search against a database of previously published alleles [31]. Sequences were aligned with MUSCLE [44] and pairwise nucleotide divergences calculated.

### Kaptive, a tool for identification of K loci in genome data

We developed an extended procedure for identification and assessment of full-length K loci among bacterial genomes based on BLAST analysis of assemblies. The procedure has been automated and is implemented in a freely available open source software tool, *Kaptive* (https://github.com/katholt/Kaptive). For full details see **Additional File 3, Figures S2 and S3.**

### Comparison of Kaptive, wzi and wzc typing results

*Kaptive, wzi* and *wzc* typing were applied to the 86 genomes that had matched serological typing information available (**Table S3, Additional File 4)**. *Wzi* alleles were determined as described above, and used to predict serotypes by comparison to the *K. pneumoniae* BIGSdb. *Wzc* sequences were extracted as above and genomes were assigned to serotypes if the sequence was <6% divergent from the corresponding reference [31].

## Results

We analysed K loci in a final dataset comprising 2503 high quality genomes, including 2298 *K. pneumoniae sensu stricto*, 144 *K. variicola*, 57 *K. quasipneumoniae* and four unclassified *Klebsiella* spp. (**Table 1** and **Table S1, Additional File 1**). Also included were 10 publicly available genomes representing the more distantly related *Klebsiella oxytoca* (**Table S4, Additional File 3**), as we hypothesised that these organisms may share capsule synthesis loci with *K. pneumoniae*. Isolates were collected between 1932 and 2014 (see **Figure S4, Additional File 3**) and from eight different geographic regions spanning six continents (see **Figure S5, Additional File 3**).

**Table 1:** *K. pneumoniae* genomes investigated in this study

| Dataset | Count | Reference | Notes |
| --- | --- | --- | --- |
| Bialek-Davenet *et al.* | 33 | [8] | Investigation of multi-drug resistant and hypervirulent clones |
| Bowers *et al.* | 160 | [49] | Isolates mostly of clonal group 258 |
| Davis *et al.* | 77 | [50] | Isolates from human UTI and animal meats in Arizona, USA |
| Deleo *et al.* | 69 | [29] | Isolates of clonal group 258 |
| Ellington | 185 | [51] | Multidrug resistant isolates from a hospital in Cambridge, UK |
| Holt *et al.* | 274 | [7] | Global diversity study |
| Lee *et al.* | 27 | [52] | Isolates from pyogenic liver abscess disease, Singapore |
| NCTC3000 | 81 | [35] | Isolates from the Public Health England NCTC reference collection |
| PATRIC | 811 | [34] | Genome assemblies submitted to GenBank |
| Stoesser *et al.* 2013 | 69 | [53] | Isolates from health-care associated infections, Oxford, UK |
| Stoesser *et al.* 2014 | 55 | [54] | Isolates collected during an outbreak in Nepal |
| Struve *et al.* | 67 | [55] | Predominantly isolates of clonal group 23 |
| The *et al.* | 76 | [3] | Isolates from two outbreaks in Nepal |
| Wand *et al.* | 35 | [56] | Historical isolate collection |
| Novel isolates | 484 | This study | Diverse collection of Australian hospital surveillance isolates |

A total of 1371 *K. pneumoniae* genomes could be putatively assigned to 63 of the 77 K-type reference loci, and a further 918 to one of the other 25 previously published K loci. Among the remaining 213 genomes, 106 were assigned to 29 novel K loci, bringing the total number of known K loci to 132. A further six genomes harboured deletion mutants of known K loci, two had IS variants of known loci and one had an IS variant of a novel locus (see below). For 93 genomes (3.7%), no K locus could be determined; however we found evidence of the presence of three or more K locus-associated genes in all such genomes and consider the lack of assignment was likely attributable to low quality sequence data (low read depth and/or fragmented assembly) rather than complete K locus deletion. Complete details of K locus assignments to all genomes are given in **Table S1** in **Additional File 1**.

A locus previously associated with K66 was identified in one *K. oxytoca* genome and the K74 locus in four *K. oxytoca* genomes; these matches were very close to the *K. pneumoniae* reference sequences (100% coverage and >93% nucleotide identity in all cases). Two novel K loci were identified in three *K. oxytoca* genomes, increasing the total known *Klebsiella* K loci to 134 (**Table S4, Additional File 3**).

We estimated the extent to which we had captured the repertoire of K locus diversity in the *K. pneumoniae* population (**Figure 1**). The rarefaction curves were estimated from (i) the full genome set for which K loci were assigned (n = 2410; grey lines in **Figure 1**); (ii) a ‘non-redundant’ genome set from which highly biased subsamples such as outbreaks were removed (n = 1081; blue lines in **Figure 1**); and (iii) genomes from the non-redundant set representing each of the distinct species, *K. pneumoniae sensu stricto, K. variicola* and *K. quasipneumoniae* (black lines in **Figure 1**). In comparison to that for the full genome set (grey), the non-redundant (blue) curve better represents the true diversity of the *K. pneumoniae sensu lato* population. Note that neither reached the total number of known K loci, since 13 of the serologically defined K loci [22] were not represented at all in our 2503 genomes. The rarefaction curves for each of the three *Klebsiella* species within the nonredundant dataset were highly similar to one another, indicating similar levels of capsule diversity within each species (**Figure 1**). There was no strong evidence of species specificity: across our entire genome collection 46 distinct K loci were identified only among *K. pneumoniae sensu stricto*, while three and two were identified only among *K. variicola* and *K. quasipneumoniae*, respectively, however these differences are likely an artefact of the much larger sample size currently available for *K. pneumoniae sensu stricto*.

**Figure 1:**
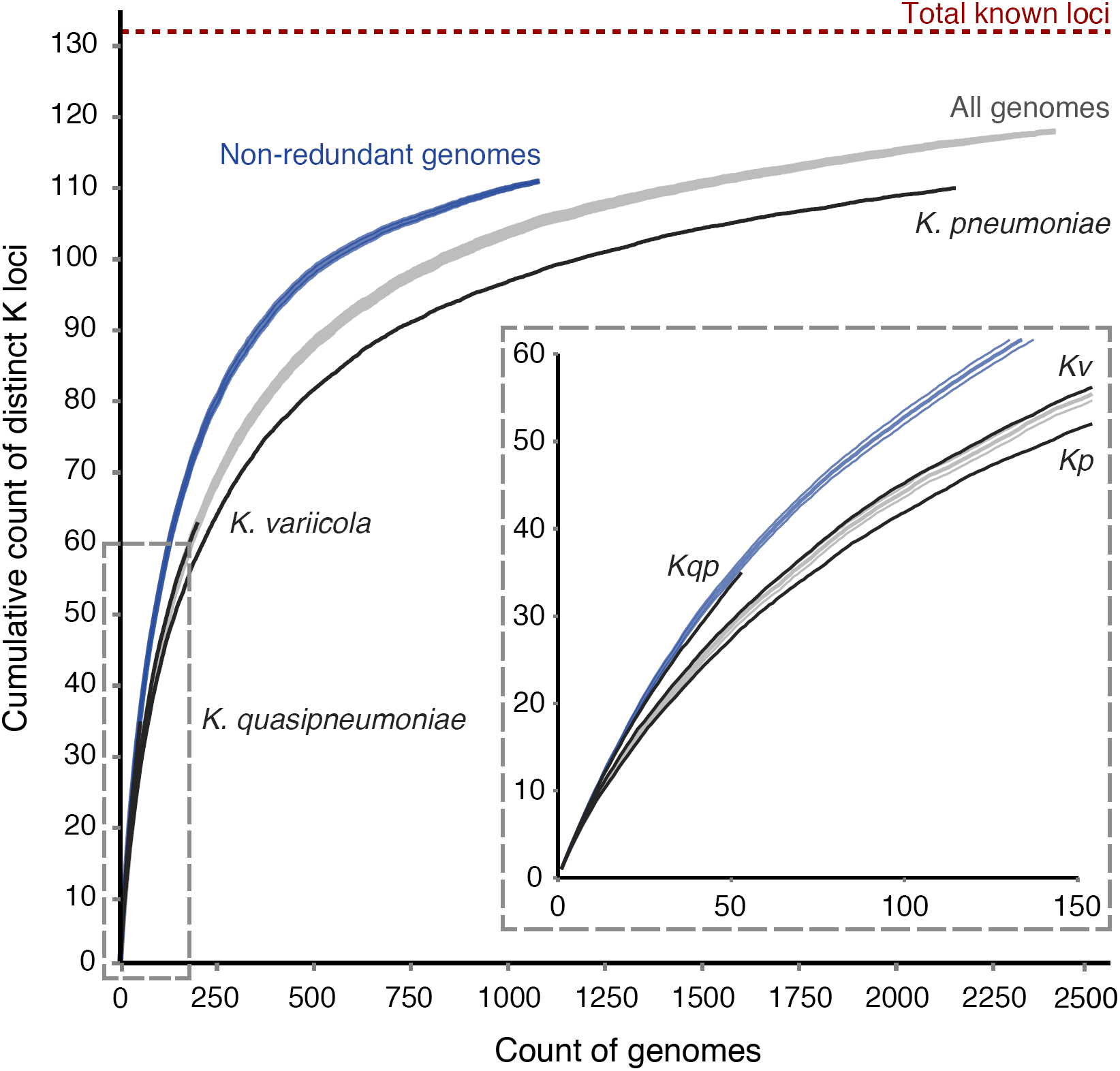
Rate of discovery of distinct K loci with increasing genome sample size. Curves indicate the accumulation of distinct K loci (mean ± SE) in different genome sets. Grey; all genomes (n=2410, excluding *K. oxytoca*). Blue; non-redundant genomes (n=1081, excludes genomes from investigations of disease outbreaks and specific clonal groups). Black; species specific genome sets (*K. pneumoniae* refers to *K. pneumoniae sensu stricto,* SE not shown). Inset box shows a zoomed view of the bottom-left section of the plot, as indicated by the dashed box.

### K locus nomenclature

We used a standardised K locus nomenclature based on that proposed for *Acinetobacter baumannii* [57]. Each distinct *Klebsiella* K locus was designated as KL (K locus) and a unique numeric identifier. The K-type reference K loci were assigned the same numeric identifier as the corresponding K-type, for example K1 is encoded by the KL1 locus. K loci for which capsule types have not yet been phenotypically defined were assigned identifiers starting from 101 (note KL101 and KL102 correspond to those previously named as KN1 and KN2).

K loci with IS insertions disrupting the region were distinguished from orthologous IS-free variants by using -1, -2. This nomenclature was consistently applied to the 10 K-type reference K loci published previously that include one or more ISs [21,22,25,26]. Deletion variants derived from a known K locus were given the suffix - D1, -D2, etc.

### K locus reference database

We curated a K locus reference database of complete annotated K locus sequences for all of the 134 loci (**Table S2, Additional File 2**). Where possible sequences were included at their full length, from the start of *galF* to the end of *ugd*. Where a previously published sequence did not span the full length of the locus or contained an IS, we substituted the complete, IS-free K locus sequence from a genome in our collection if available (39 of 51). Where no naturally occurring IS-free variants were available, we manually generated an IS-free synthetic sequence (**Table S2, Additional File 2**). IS-free sequences are included in a primary K locus reference database, while all available IS or deletion variant K locus references are included in an accompanying variant database, both available at https://github.com/katholt/Kaptive.

### K locus structures

All of the novel K loci identified in this study conformed to the common structure described previously (**Figure S6**) [20–22]. We also identified six deletion variants within our genome collection (KL5-D1, KL20-D1, KL30-D1, KL62-D1, KL106-D1 and KL107-D1). Each putative deletion variant was missing several common genes but the remaining regions showed a high degree of similarity to other apparently complete K loci, which we suggest represent the ancestral forms. Isolate NCTC10004 (recorded as serotype K11 in the UK National Culture Type Collection) and four other genomes carried a K locus that was nearly identical to the previously published K11 K locus reference sequence [22]. However, the latter lacked the essential *wzx* gene plus two other neighbouring genes, and was not identified among any other genomes. We assume the NCTC10004 locus represents the full length KL11 locus and designate the original K11 reference as KL11-D1 (note it is unclear whether the original sequenced reference isolate had retained the ability to produce a capsule, since the serological typing was performed decades earlier [22]). In four of the deletion variants the deleted region was replaced by an IS, which may have mediated the deletion.

Of the other IS related variants, KL157-1 contained an IS903 family IS without an obvious deletion. In addition, we identified two novel IS variants of K-type reference loci (KL15-1 and KL22-1), and five IS-free variants of K-type reference loci (KL3, KL6, KL38, KL57, KL81), plus one other previously published K locus (KL103). In total, seven IS insertions were associated with neither a deletion nor a rearrangement event. In contrast, the KL22-1 locus included a translocation of part of the lipopolysaccharide (LPS) locus to the centre of the K locus, plus an inversion of the 3’ portion of the K locus (**Figure 2**). The translocated and inverted regions were bound at each end by a copy of *ISKpn26*.

**Figure 2:**
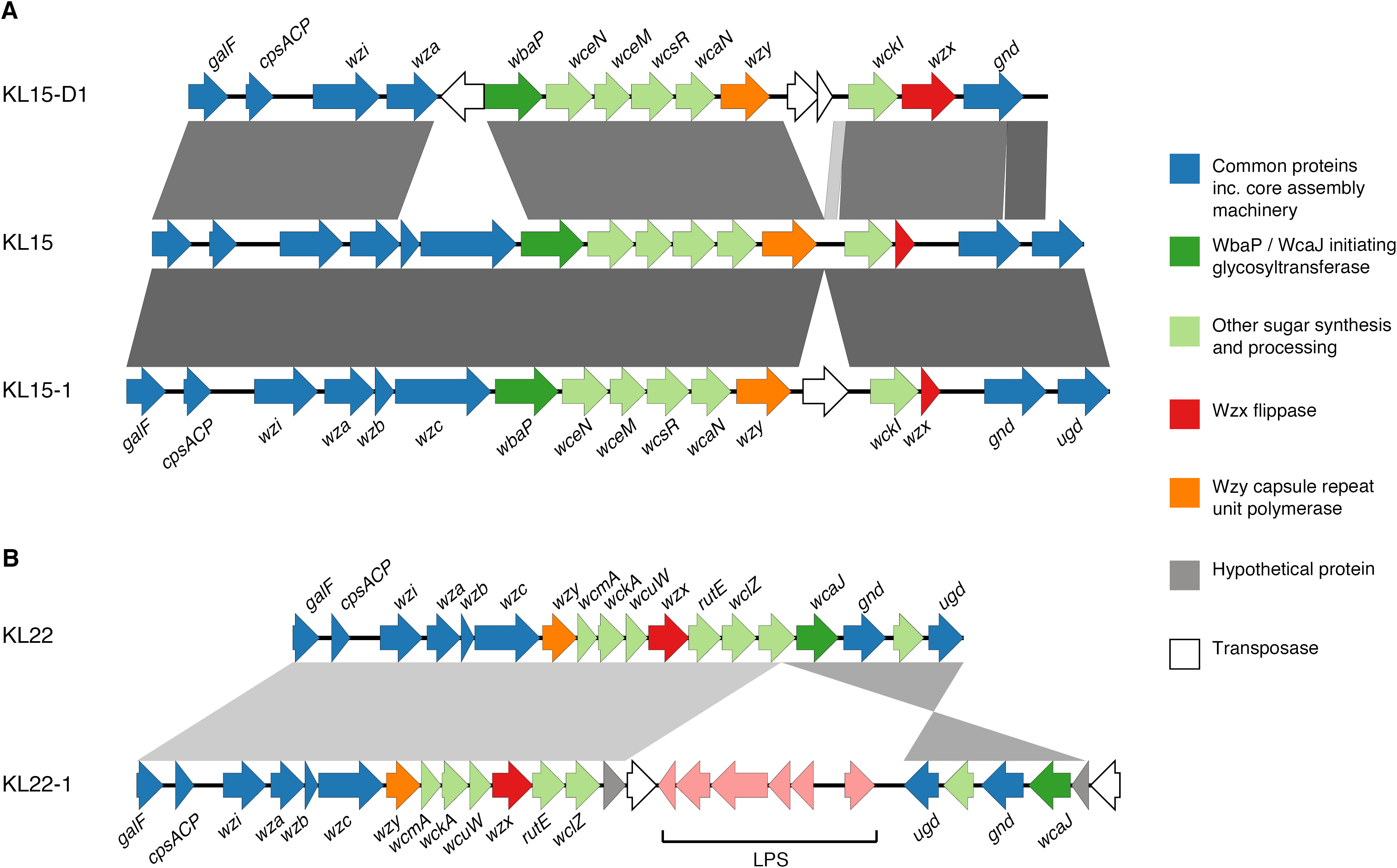
Example K locus structures and comparisons Coding sequences are represented as arrows coloured by predicted function of the protein product and labelled with gene names where known. Grey bars indicate regions of similarity identified by BLAST comparison, darker shading indicates higher sequence identity. **(A)** Comparison of deletion variant KL15-D1 and insertion sequence variant KL15-1 with the synthetic KL15 locus. **(B)** Comparison of the insertion sequence variant KL22-1 with the K-type reference KL22 locus. The downstream LPS (lipopolysaccharide synthesis) operon (pink arrows) has been translocated into the K locus.

ClonalFrameML analysis of the nucleotide sequences of common K locus genes identified a high number of putative recombination events (n=382) between the reference K loci. These events were not distributed equally across the nucleotide alignment (**Figure 3A**); rather the genes closest to the central variable region of the K locus were affected by a greater number of recombination events compared to those at the ends of the locus.

**Figure 3:**
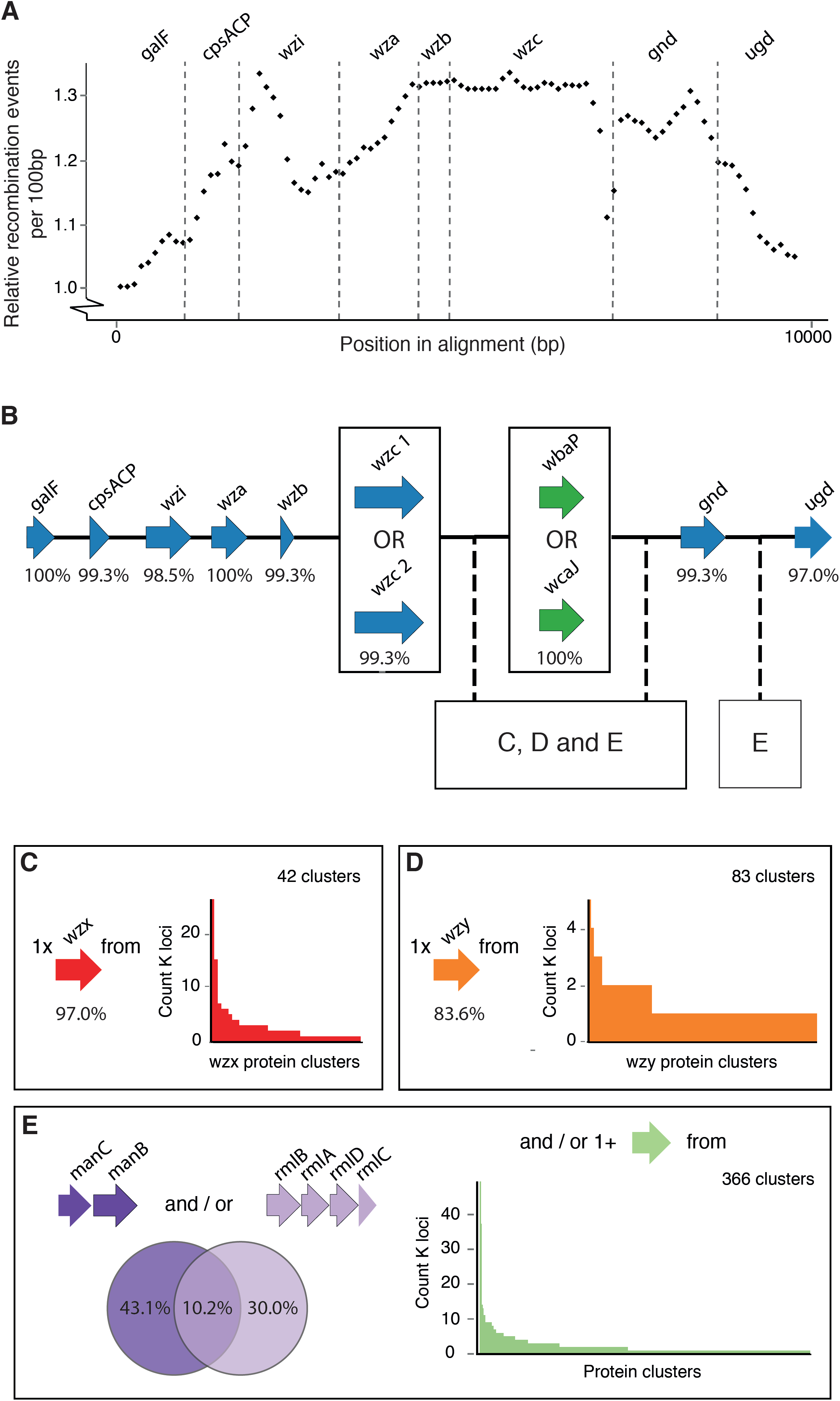
Composition and diversity of *Klebsiella* K loci **(A)** Putative recombination events among the common K locus genes. Values plotted are relative number of events per 100 bp window, inferred using ClonalFrameML. **(B)** Representation of a generalised K locus structure. Arrows represent K locus coding regions coloured by predicted protein product as in **Figure 2**. Percentage values indicate the number of reference K loci containing each gene (total 134 references). Note that 13 of the K locus references partially or completely exclude *ugd,* although it is known to be present in 11 of these loci [22]. Thus we counted these 11 as containing *ugd.* The locations within this structure at which *wzx* (C), *wzy* (D) and sugar processing genes (E) have been found to occur are indicated. **(C, D, E)** Diversity of proteins encoded by *wzx* (C), *wzy* (D) and sugar processing genes (E) annotated amongst the 134 K locus reference sequences. Bar charts indicate the frequency of each predicted protein cluster.

### Variation in K locus gene content

A total of 2675 predicted proteins from 134 complete K loci were clustered at various identity levels (see **Methods**), resulting in 1496 to 508 clusters. As the identity threshold was reduced the number of clusters continued to fall and showed no signs of stabilising, even between 50% and 40% identity (**Figure S7**), and we believe the latter is a lower bound for sensible comparison. At 40% identity, the core capsule assembly proteins GalF, Wzi, Wza, Wzb, Gnd and Ugd each formed a single cluster, and were present in nearly all loci (**Figure 3**). The Wzc sequences clustered into two groups, and each locus encoded one Wzc protein (except KL50). In contrast, Wzx (flippase) clustered into 42 groups and Wzy (capsule repeat unit polymerase) clustered into 83 groups, highlighting the extreme diversity of these proteins compared to the other core capsule assembly machinery proteins (**Figure 3**).

There were 374 clusters among the remaining proteins, almost all of which were associated with sugar synthesis and processing (**Figure 3**). The initiating sugar transferase proteins WbaP (undecaprenyl-phosphate galactosephosphotransferase) and WcaJ (undecaprenyl-phosphate glucose phosphotransferase) were grouped into two clusters. These proteins are considered essential for capsule synthesis. Concordantly each locus encoded a single protein from one of these two clusters. RmlB, RmlA, RmlD and RmlC, which are associated with synthesis and processing of rhamnose and typically encoded together in a single operon, were each represented by a single cluster. Similarly, the mannose synthesis and processing proteins, ManC and ManB, were grouped into a single cluster each. The associated operons *rmlBADC* and *manCB* were present in 55 and 73 K loci, respectively (14 loci contained both operons, **Figure 3**). In contrast, 360 of the remaining 366 protein clusters were present in fewer than ten K loci each (**Figure 3E**).

Co-occurrence analysis identified 18 correlated groups of K locus proteins, ranging in size from two to five protein clusters (pairwise Jaccard similarity ≥0.61 for all pairs in the group, **Figure 4** and **Table S5, Additional File 5**). One group included the four Rml protein clusters; interestingly this group also included a WcaA glycosyltransferase, which was present in 67.3% of *rm/BADC-containing* K loci and no rm/BADC-negative loci (Χ-squared = 70.09, p-value < 2.2e-16 by two-sided proportion test). Similarly another group included the ManCB proteins and the putative mannosyl transferase, WbaZ, which was present in 65.8% of manCB-containing K loci and one *manCB-* negative locus (Χ-squared = 56.159, p-value = 6.683e-14 by two-sided proportion test). In addition, several groups included proteins for which the associated genes were located sequentially in their K loci (e.g. *wckG, wckH* and *wzx* in KL12, KL29 and KL42) consistent with linked gene transfer.

**Figure 4:**
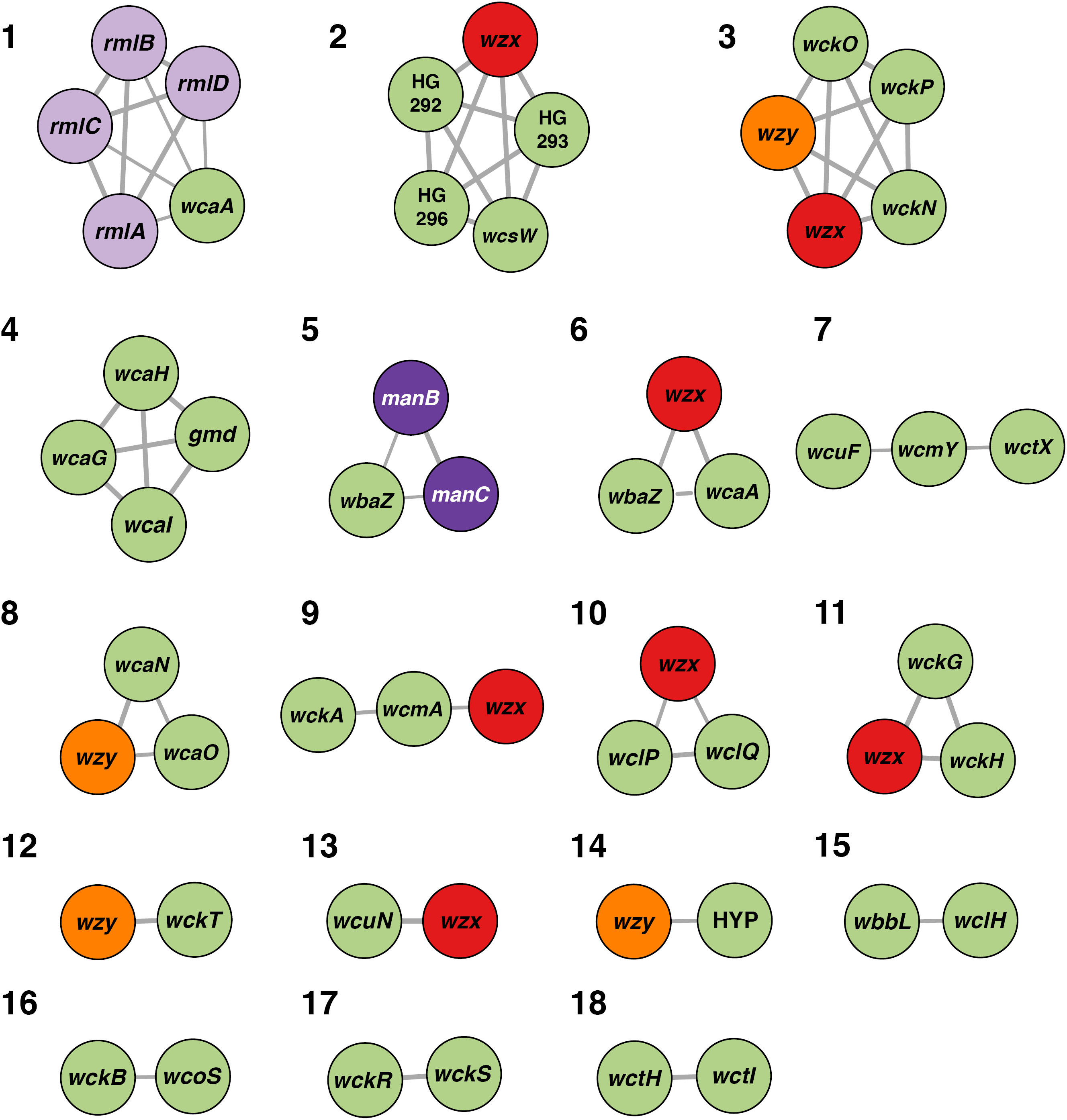
Co-occurrence of *wzx, wzy* and sugar processing genes across reference K loci Nodes represent genes, labelled by name (HYP: hypothetical protein) and coloured by protein product as in Figures 2 and 3: Wzx (red), Wzy (Orange), mannose synthesis/processing proteins (dark purple), rhamnose synthesis/processing proteins (light purple), other proteins (green). Edge widths are proportional to Jaccard index (7) and are shown for all pairs where *J* ≥0.61. Numbers represent co-occurrence group assignments as defined in **Table S5, Additional File 5**.

### Diversity of wzc and wzi gene sequences

We confidently assigned *wzi* alleles to 2461 *K. pneumoniae* genomes, including 390 distinct alleles, 218 of which were novel. Median pairwise nucleotide divergence was 7%. Among the non-redundant genome set there were 54 *wzi* alleles represented by at least five genomes and of these, 15 (28%) were associated with more than one K locus type (**Table S1, Additional File 1**). Much of the *wzi* allelic variation appeared to result from accumulation of mutations within K loci. Among the 67 K loci for which we had ≥5 representative sequences, 64 (95.5%) were associated with two or more *wzi* alleles, and there was a general trend towards increasing *wzi* allelic diversity with increasing K locus representation (**Figure 5**).

**Figure 5:**
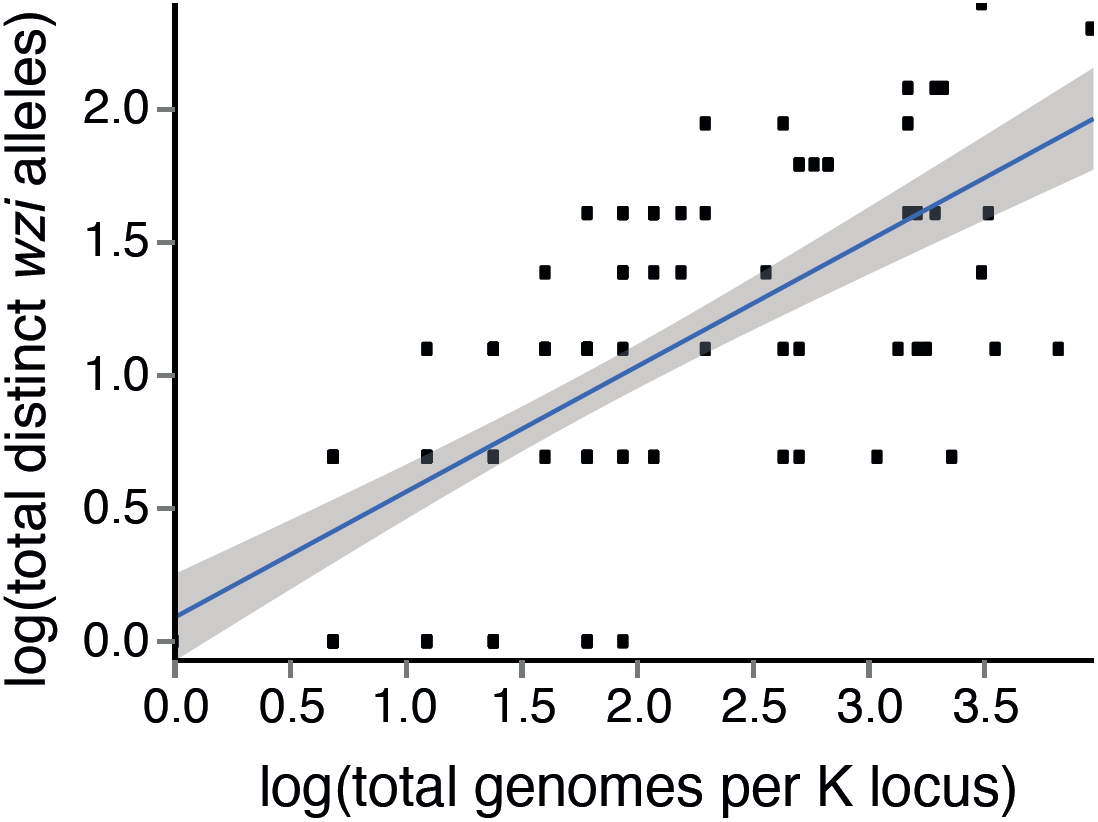
Allelic diversity of *wzi* Within K locus *wzi* allelic diversity increases with total K locus representation. The blue line represents the least-squares regression and grey shading indicates the 95% confidence interval.

We extracted *wzc* sequences from 1041 of 1082 genomes in the non-redundant genomes set (**Figure 6**). In general, genomes sharing the same K locus (n=6262 pairwise observations) showed lower *wzc* nucleotide divergence than those with different K loci (n=491,775 pairwise observations), but the distributions overlapped substantially (**Figure 6**). Notably, there were five distinct combinations of K loci for which one or more pairs harboured *wzc* sequences that were <6% divergent (the cut-off for K-type assignment as described in [31]; KL1 and KL112, KL9 and KL45, KL15 and KL52, KL30 and KL104, KL40 and KL135). Conversely, some K loci (KL45, KL112) had more than 25% *wzc* nucleotide diversity between representatives of the same K locus.

**Figure 6:**
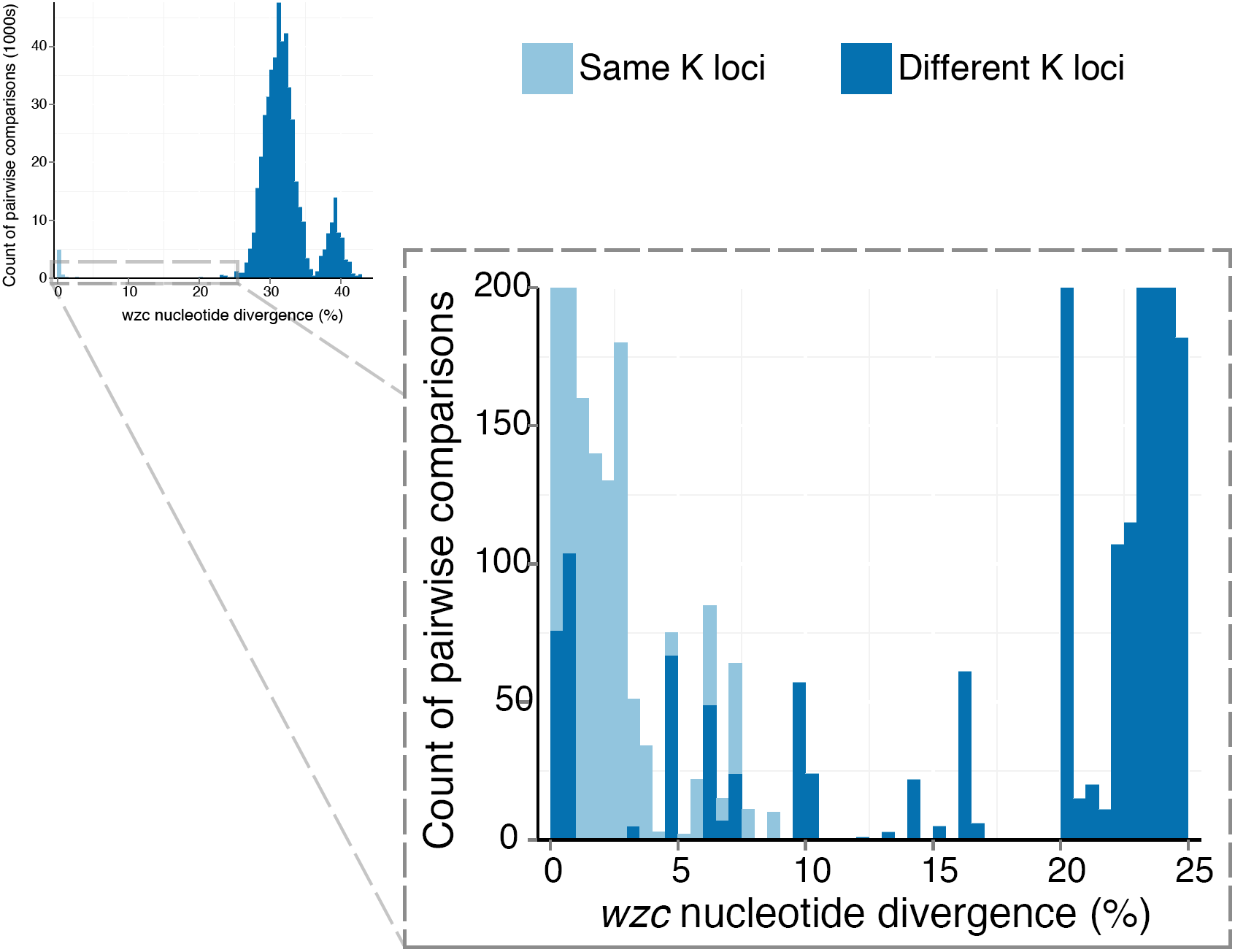
*wzc* nucleotide diversity Barplots showing distribution of pairwise *wzc* nucleotide divergence for pairs of genomes with the same (light blue) or different (dark blue) K loci. The inset box shows a zoomed view of the lower end of the distribution.

### Kaptive - capsule locus (K locus) typing and variant evaluation from genome data

To facilitate easy identification of K loci from genome assemblies, we developed the command-line software tool *Kaptive*, which is an extension of the analysis procedure described above, as shown in **Figure 7** (also see **Additional File 3**). We used *Kaptive* with our primary *Klebsiella* K locus reference database to rapidly type the K loci in our collection of 2503 *Klebsiella* genomes, and obtained confident K locus calls for 2412 genomes (96.4%, see **Additional File 3** for further details).

**Figure 7.**
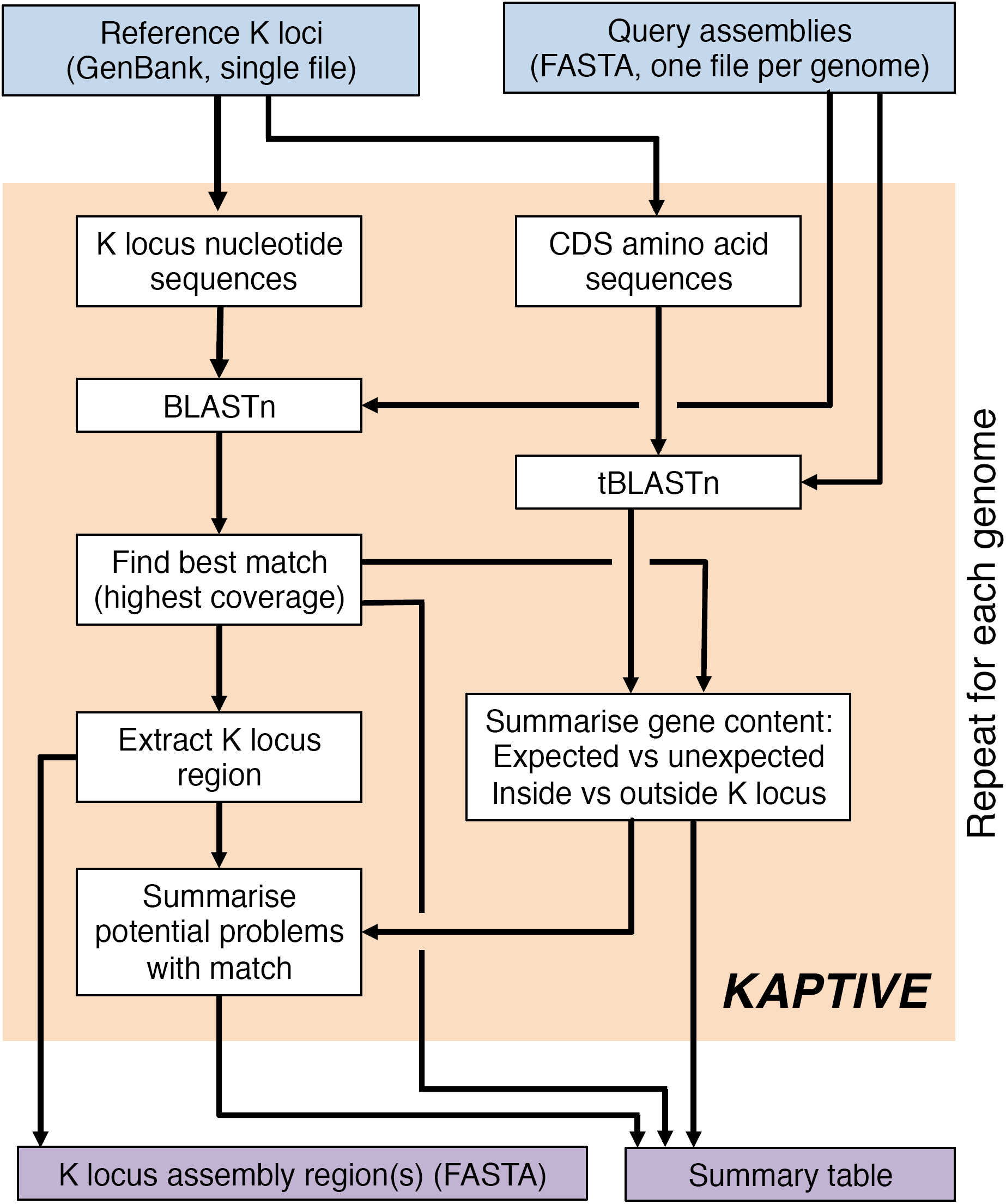
Summary of the *Kaptive* analysis procedure *Kaptive* takes as input a set of annotated reference K loci in GenBank format and one or more genome assemblies, each as a single FASTA file of contigs. *Kaptive* performs a series of BLAST searches to identify the best-match K locus in the query genome and assess the presence of genes annotated in the best-match locus (expected genes) and those annotated in other loci (unexpected genes) both within and outside the putative K locus region of the query assembly. The output is a FASTA file containing the nucleotide sequence(s) of the K locus region(s) for each query assembly and a table summarising the best-match locus, gene content and potential problems with the match (e.g. the assembly K locus region is fragmented, expected genes are missing from the K locus region or at low identity, or unexpected genes are present) for each query assembly. CDS = coding sequence.

We compared the K locus calls from *Kaptive, wzc* and *wzi* typing to serological typing results for 86 isolates for which both genome and serology data were available ([7,18,35], see **Table S3, Additional File 4**). Five of six isolates that were non-typeable by serological techniques were identified by *Kaptive* as carrying KL16, KL54, KL81, KL111 and KL149. The KL16, KL54 and KL81 calls were in agreement with *wzc* and *wzi* typing results; the other two K loci were not present in the *wzi* or *wzc* schemes and so were not typeable by those methods. Among the 80 serologically typeable isolates, the three molecular methods were generally in agreement with one another, although concordance with recorded phenotypes was quite low (6574%, Table S3). Call rates were highest for *Kaptive* (95%), followed by *wzc* (89%) and *wzi* typing (75).

## Discussion

The number of distinct *Klebsiella* K loci (now 134) is striking and exceeds that described for capsule synthesis loci in other bacterial species such as *A. baumannii, Streptococcus pneumoniae* and *Neisseria meningitidis*. Furthermore, the diversity is an order of magnitude greater than that recently described for *Klebsiella* LPS, the other major *Klebsiella* surface antigen [28]. This suggests that the K locus is subject to strong diversifying selection. Given that these bacteria are not obligate pathogens and are ubiquitous in non-host-associated environments [58,59], it seems likely that the factors driving selection are not immune pressures from humans or other hosts, but may include phage and/or protist predation.

Two novel K loci were identified from *K. oxytoca*, a close relative of *K. pneumoniae*. The KL66 and KL74 K-type reference loci were also identified among *K. oxytoca* genomes. Little is known about *K. oxytoca* capsules, though a report from Japan in 2012 identified several other *K*. pneumoniae-associated capsules among *K. oxytoca* isolates from blood and bile infections [60]. Together these findings indicate that *K. oxytoca* is able to exchange genetic material with *K. pneumoniae*. Therefore, *K. oxytoca* represents a potential reservoir of virulence, drug resistance and other genes for *K. pneumoniae*, and warrants greater research attention.

Our analysis confirms there are strong constraints on the structure of K loci, which generally include *galF, cpsACP, wzi, wza, wzb* and *wzc* at the 5’ end, *gnd* and *ugd* at the 3’ end, and a highly variable set of genes in the centre (Figure 3). Our data also reveal the extensive diversity of proteins encoded in the variable central region, with 499 unique proteins identified across the 134 K loci. These genes ranged in frequency from 0.7% to 54.7% of the K loci. Among those represented in at least three K loci, approximately half co-occurred in groups ranging from two to five genes.

The molecular evolutionary events driving K locus diversification are not yet well understood, but likely include a combination of point mutation, IS-mediated rearrangements, and homologous recombination within the locus, resulting in the mosaic structure summarised in Figure 3. It has been shown, in both *A. baumannii* [61,62] and *S. pneumoniae* [63], that recombination within the capsule synthesis locus can drive capsule exchange between distinct clones. We recently speculated that this may also be true for *K. pneumoniae* [27] and the recombination analysis presented here supports this theory. The genes closest to the central variable region of the K locus (i.e. *wzb, wzc* and *gnd*), showed evidence of the greatest number of recombination events, consistent with the hypothesis that they act as regions of homology for recombination events that shuffle the central region of the locus.

The prediction of capsule phenotypes from genome data is complex, as capsule expression is a highly regulated process that involves loci outside the K locus region [64], and so presence of an identical K locus sequence does not guarantee an identical phenotype. However it is likely that K loci encoding distinct sets of proteins are associated with distinct capsule phenotypes, as is the case for the vast majority of K-type reference strains [22]. Therefore, our data suggest that there are at least 134 distinct *Klebsiella* capsule types. Note that this is a lower bound estimate since there are likely additional K loci in the wider population that were not in the current genome collection, and our analysis did not capture differences that may arise from point mutations and small-scale insertions or deletions (e.g. in the case of K22 and K37 described previously [22]). Furthermore, while we did not attempt to thoroughly characterise IS variants, several such variants were apparent. The potential functional impacts of IS insertions likely vary depending on their location in the locus, but may include up-regulation, loss of capsule production and/or more subtle changes in sugar structures [65–67]. However, functional studies are required to understand these effects and to improve the prediction of phenotypes.

Serological typing of *Klebsiella* isolates is notoriously difficult and rarely performed. We were able to compare genotypes (whole-locus typing using *Kaptive*, as well as *wzi* and *wzc* typing schemes) with phenotypes on just 86 isolates for which both sequences and serotypes were available. Of the 19 discordant genotype vs phenotype results, two were due to deletion variants and were resolved by running *Kaptive* with the K locus variants database. Interestingly, one of these isolates was non-typeable by serology, *wzi* or *wzc* typing, but recognized as a specific K locus deletion variant by *Kaptive*. This highlights a benefit of our whole-locus typing approach; it provides epidemiologically relevant information even when the K locus is interrupted. Another isolate was serologically typed as K54 but genotyped by *Kaptive* as KL113, which has sequence homology with KL54 (>84% nucleotide identity over 76% of the locus) and may encode a serologically similar or crossreacting capsule. The other cases of discordance had no obvious explanation, however it is likely that some result from serological typing errors or from mutations arising during subculture (as identified for the K11 reference isolate above), neither of which we were able to check. Some discordance may also be due to unpredictable serological cross-reactions.

Given the problems with serotyping and the comparative robustness and widespread access to genome sequencing, we anticipate that genotyping will remain the preferred method for tracking capsular diversity in *Klebsiella*. Due to the extensive diversity and potential for ongoing evolution, we strongly advocate for classification based on complete, or near complete K locus sequences, rather than single genes such as *wzi* or *wzc*, which can be misled by substitutions and horizontal gene transfer. *Kaptive* analyses the full-length K locus nucleotide sequences and assesses the presence of all K locus associated genes by protein BLAST search, thus the approach is resilient to spurious results that may arise due to sequence divergence. Furthermore, the information provided allows users to determine confidence in the results and to identify putative novel K loci or variants of known loci if desired (see **Additional File 2**).

## Conclusions

We report an investigation of K loci among a large collection of 2513 *Klebsiella* genomes. We identified 31 novel K loci, increasing the total number of known loci to 134, almost twice the number of serologically defined K-types. We defined a standardised *Klebsiella* K locus nomenclature and developed a curated reference database, which captures the majority of the extensive diversity in the *K. pneumoniae* population. Lastly, we developed a simple program, *Kaptive*, for the detection of reference K loci from genome assemblies. These new resources will greatly facilitate evolutionary investigations and genomic surveillance efforts for this and other important bacterial pathogens.

## List of Abbreviations

K-type: capsule type
K locus: capsule synthesis locus
IS: insertion sequence
CDS: coding sequence
LPS: lipopolysaccharide

## Additional Files

### Additional file 1 (Excel spreadsheet, xlsx)

**Table S1.** *K. pneumoniae* genome data analysed in this study. Accession numbers, K locus designations and summarised *Kaptive* typing results are provided.

### Additional file 2 (Excel spreadsheet, xlsx)

**Table S2.** *Klebsiella* K locus references, accession numbers and isolate names.

### Additional file 3 (Word document, docx)

**Supplementary Methods and Results**

**Table S4.** *K. oxytoca* genomes analysed in this study and K locus designations.

**Figures S1 - S7.**

### Additional file 4 (Excel spreadsheet, xlsx)

**Table S3.** *Kaptive, wzi, wzc* and serological typing results for 86 *K. pneumoniae* genomes for which serological typing information were available.

### Additional file 5 (Excel spreadsheet, xlsx)

**Table S5.** Jaccard similarity scores for 115 K locus protein clusters for which the associated genes were present in at least three K loci.

## Declarations

### Availability of data and material

The datasets generated and/or analysed during the current study are available in the European Nucleotide Archive and/or PATRIC genome database, accession numbers are listed in **Table S1**. Novel K locus nucleotide sequences have been deposited in GenBank (accessions listed in **Table S2**) and are also distributed together with our reannotated and curated set of all known K loci at https://github.com/katholt/Kaptive.

## Authors’ Contributions

KLW and KEH designed the study, collected and analysed data, designed the *Kaptive* tool and wrote the manuscript. RRW designed and implemented the *Kaptive* tool and wrote the manuscript. CG and AJ generated serological and sequence data for Australian isolates. RF and NT contributed to data analysis and interpretation. All authors read and approved the final manuscript.

